## Supplementary Information for "The molecular pH-response mechanism of the plant light-stress sensor PsbS"

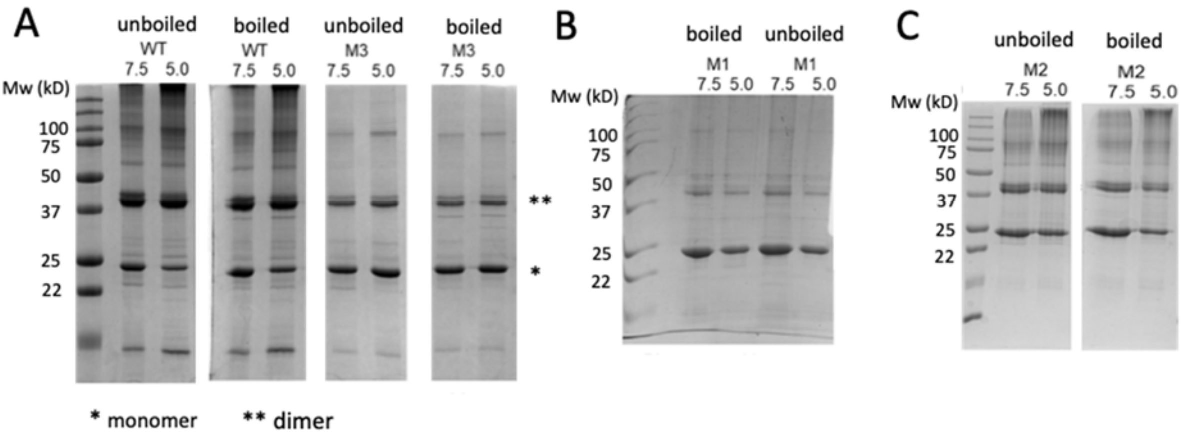


**Figure S1. SDS-page gel images of PsbS at pH 7.5 and pH 5.0 conditions. A: Wild-type PsbS and E71Q/E176Q mutant M1/M2; B: E71Q mutant M1 and C: E176Q mutant M2.**

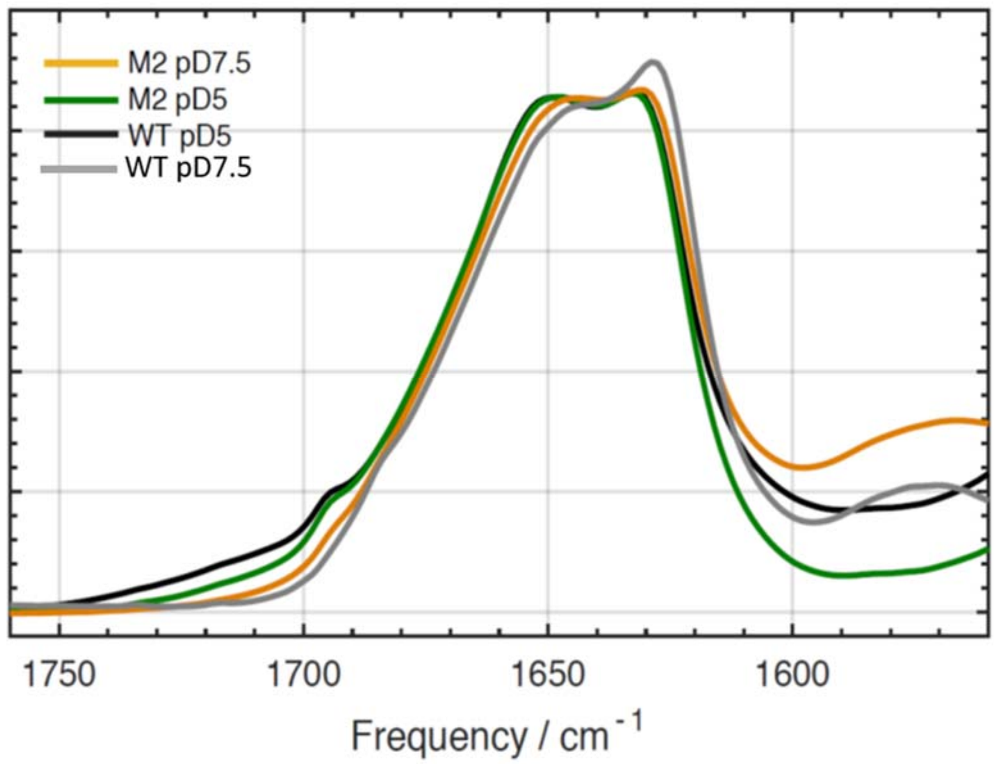


**Fig. S2. Overlaid FTIR spectra of wild-type *P. patens* PsbS and the M2 mutant (E176Q) at pD 7.5 and pD 5.0. The spectra were reproduced from Fig. 4A,C and scaled according to their integrated Amide I absorption between 1610 and 1690 cm$^{-1}$.**

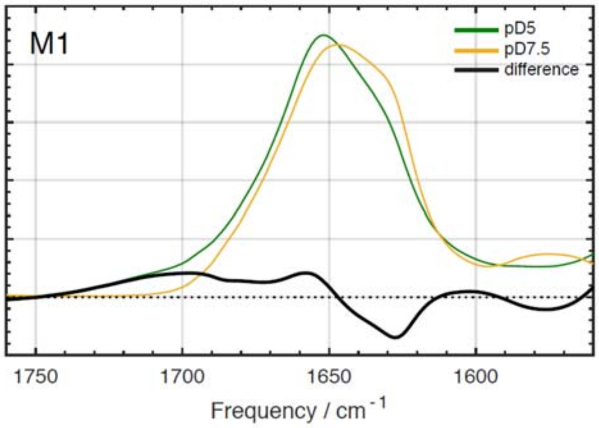


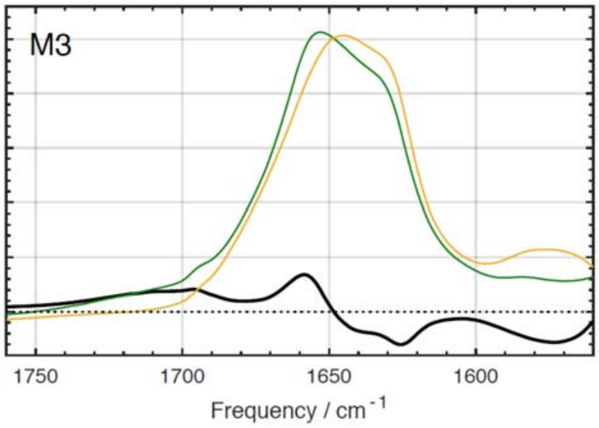


**Fig. S3. FTIR spectra of A: the M1 (E71Q) mutant; B: the M1/M2 (E71Q / E176Q) mutant at pD 7.5 (orange) and pD 5.0 (green) and the difference spectra of pD 5.0 minus pD 7.5 (black). All spectra were taken in $D_2O$ buffer.** The Amide I spectra of M1 and M1/M2 mutant do not resemble the wild type spectra at either pD condition**s** (Fig. 3A). The luminal loop containing Glu-1 contains an additional conserved, second Glu close to TM2 (E76 for spinach PsbS and E78 for *P. patens* PsbS), and, for *P. patens* PsbS, also contains an Asp residue (D69). At neutral pH, those residues will have a negative charge, while in M1 and M1/M2 the Glu-1 site is neutralized by Gln replacement, resulting in a charge distribution along the luminal loop stretch that neither reflect the luminal loop charges of wild-type PsbS at neutral pH, nor reflects the loop charges of the wild type at low pH conditions. This may lead to a non-native fold of the luminal loop stretch in M1 and M1/M2, accounting for the difference in secondary structure compared to the WT. The difference spectra of M1 and M1/M2 in the Amide I region show a well-resolved positive band at 1660 $cm^{-1}$ and negative signal intensities between 1625-

1640 cm$^{-1}$. The difference spectrum of M1/M2 shows that if both active Glu are mutated, conformational changes occur that are not present in the wild type and that must be induced by protonation of other protonable residues at low pH. Indeed, both the IR and NMR data indicate that going from pH 7.5 to pH 5.0, all the protonable Asp and Glu residues undergo a change in protonation state. However, given that the secondary structures of the M1 and M1/M2 mutants significantly differ from the wild-type structure, these changes are likely unrelated to the native PsbS response to pH.

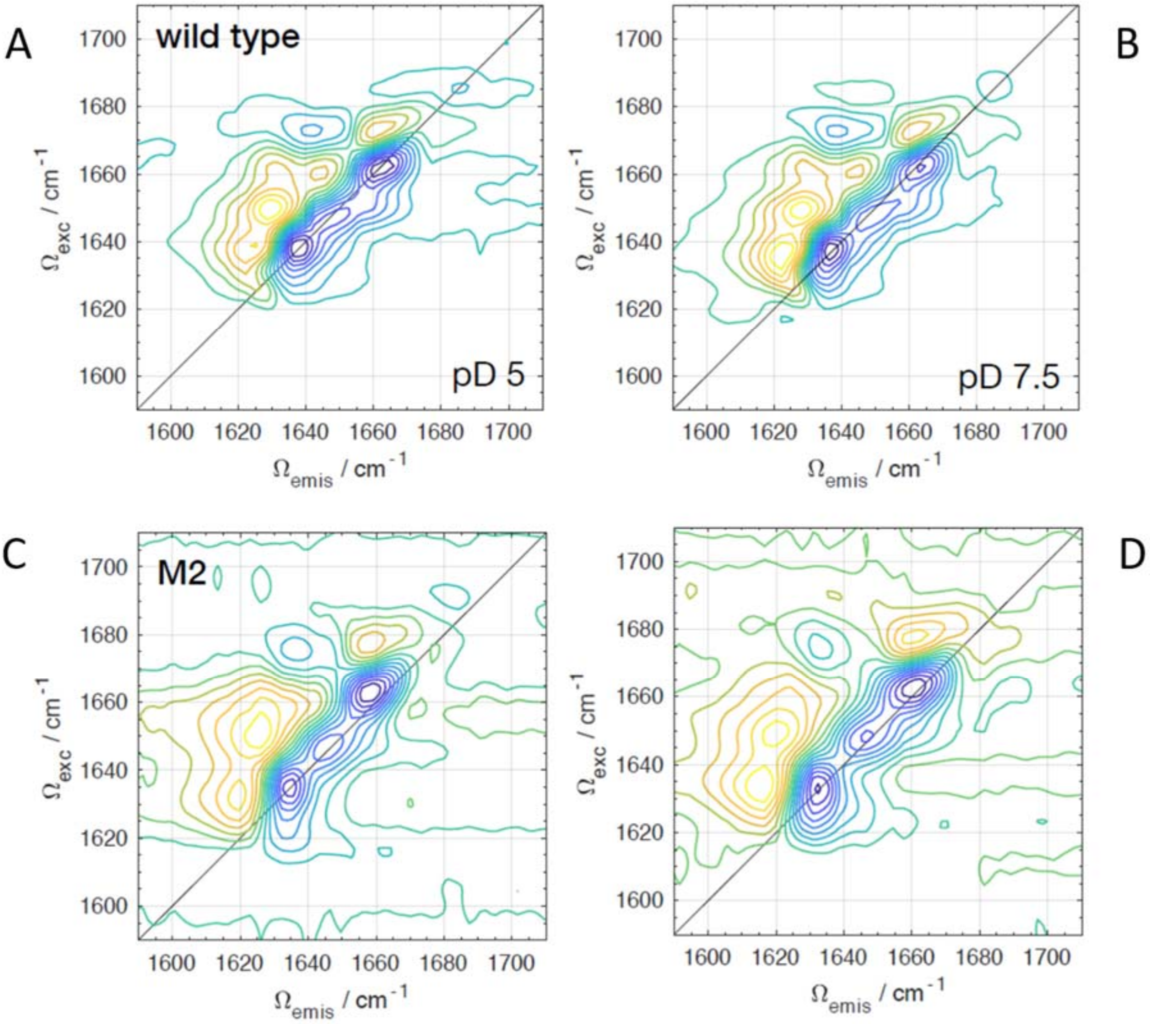


**Fig. S4. Steady-state 2DIR correlation spectrum of PsbS of (A) wild type in pD 5.0 buffer; (B) wild type in pD 7.5 buffer; (C) the M2 mutant in pD 5.0 buffer; (D) the M2 mutant in pD 7.5 buffer.**

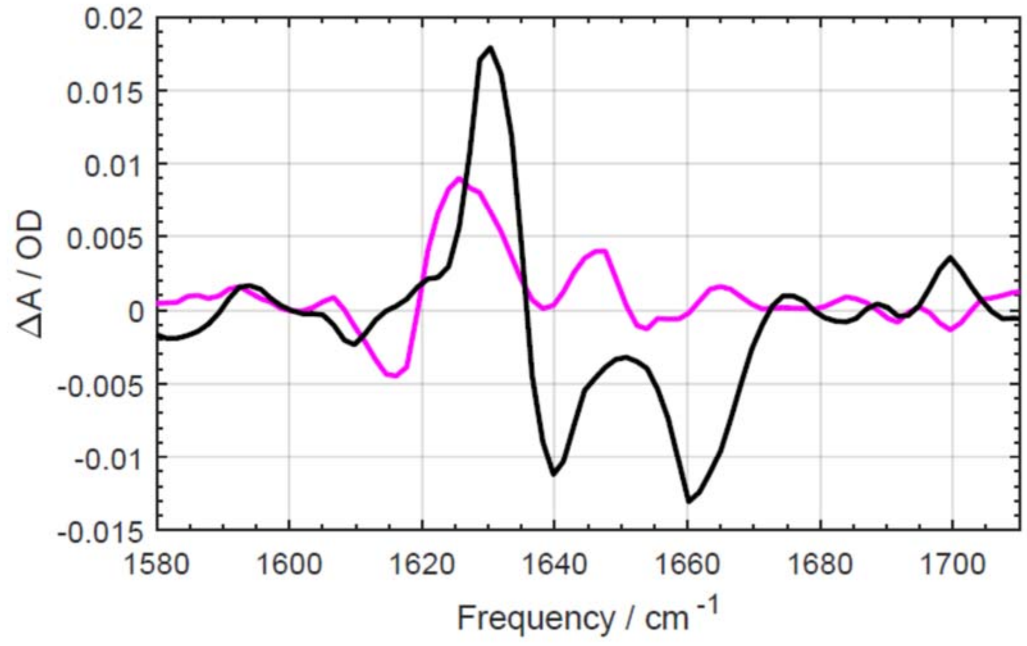


**Figure S5: diagonal slices of wild type (black line) and M2 (magenta line) 2DIR difference spectra to indicate amplitudes of 1630, 1638 and 1660 cm$^{-1}$ bands, taken from the data shown in Fig. 4B and 4C.**

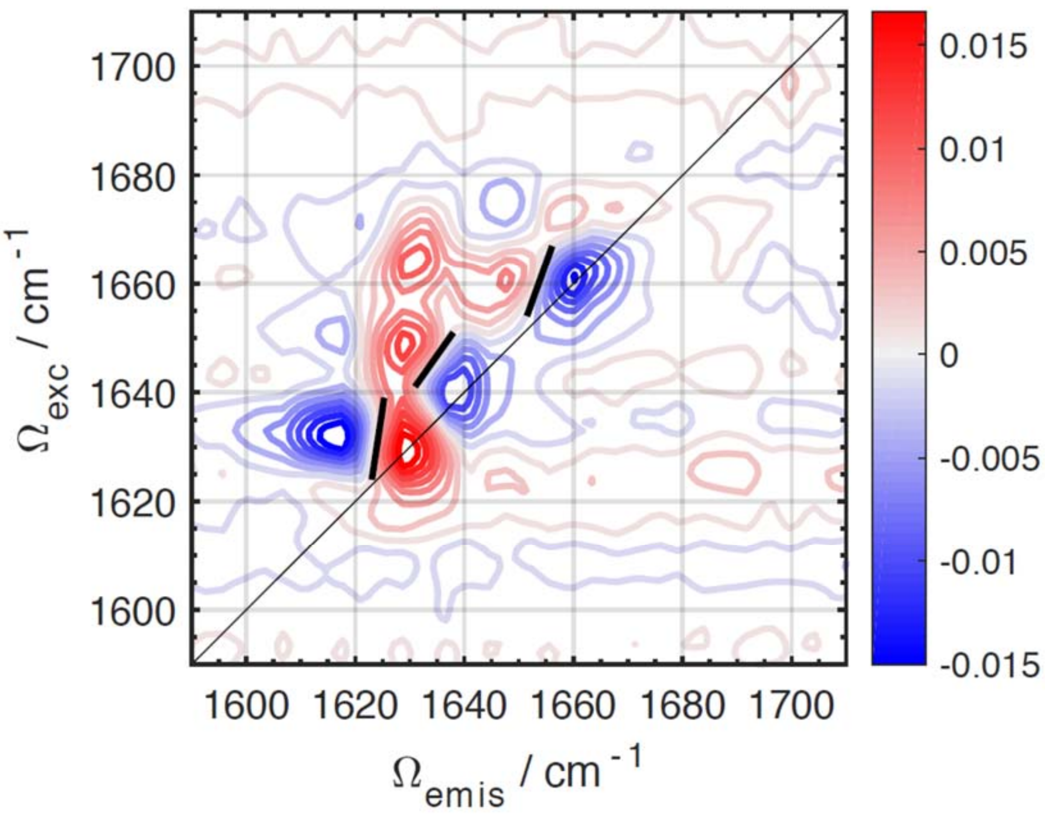


**Fig. S6. Illustration of nodal line slopes (NLS) for the diagonal peaks (thick black lines), calculated by applying a linear fit to the zero crossing (nodal line) within the FWHM of each peak (1). Calculated values are (in degrees from the diagonal): 1630 peak - 36.4; 1638 peak - 10.0; 1660 peak - 23.4. The 2DIR difference spectrum was reproduced from Fig. 4B.**

1. Kwac K & Cho MH (2003) Molecular dynamics simulation study of N-methylacetamide in water. II. Two-dimensional infrared pump-probe spectra. *Journal of Chemical Physics* 119(4):2256-2263.
